# Indicators to assess interannual variability in marine connectivity processes: a semi-theoretical approach

**DOI:** 10.1101/2024.01.12.575479

**Authors:** Morane Clavel-Henry, Nixon Bahamon, Jacopo Aguzzi, Joan Navarro, Miguel López, Joan B. Company

**Affiliations:** Institut de Ciències del Mar, Consejo Superior de Investigaciones científicas, Barcelona, Spain; Institut Català de Recerca per a la Governança del Mar, Barcelona, Spain

**Author notes:** Corresponding author (MCH). Open Research statement: A code is provided as private-for-peer review via the following link: https://figshare.com/s/a4f26cf6e3148934877e. The code will be permanently archived and made public in the Figshare repository (with a doi) if the paper is accepted for publication.

## Abstract

Oceanographic connectivity in an effective network of protected areas is crucial for restoring and stabilising marine populations. However, temporal variability of connectivity is rarely considered as a criterion in designing and evaluating marine conservation planning. In this study, indicators were defined to characterise occurrence, strength and frequency of the temporal variability in connectivity in a northwestern Mediterranean Sea area. Indicators were tested on semi-theoretically-estimated connections provided by the runs of a passive particle transport model in a climatological year and in three years between 2006 - 2020, showing large deviation from the climatological year. The indicators compared the temporal variability in connectivity of four zones, highlighting differences in connectivity due to their locations and the mesoscale hydrodynamics, and identifying areas that require further investigation. The three indicators also showed that the temporal variability in connectivity was influenced by the duration and depth of particle transport, although no consistent pattern was observed in the indicator variations of the compared zones. Provided that specific objectives will be given when parameterising transport models (i.e., selection of focus species and time period), indicators of temporal variability in connectivity have potential to support, and correctly implement, spatial conservation planning, prioritise the protection of marine resources, and measure the effectiveness of Marine Protected Areas, in line with a long-term vision of ocean management.

## Introduction

Oceanographic connectivity can be broadly defined as the existence of “a specific path between identified places” [1]. This definition relies heavily on the hydrodynamics that govern the transport of elements in the oceans. Oceanographic connectivity is associated with inert elements (e.g., pollutants, microplastics), and living elements (e.g., fish larvae), which are exposed to seasonal variability of the hydrodynamics and to the variation of the hydrodynamic fields with depth. Many elements are only transported by the marine currents, with some of them, having a limited control over their transportation like early life stages of many marine species. Other elements have complete control over the currents and dominate their dispersal in the oceans like, for example, highly migratory species. For marine beings, connectivity involves and impacts diverse biological scales (i.e., individuals, species, populations, communities, and ecosystems). Connectivity is fundamental to guarantee gene flow and the spatial structure and dynamics of populations within the network and subject to extreme oceanic events [2, 3]. The genetic mixing brought by the newly arrived individuals contributes to the stability and resilience of the population against environmental shifts and pathogen aggressions [4]. Besides, marine connectivity events may produce impactful changes in the ecological status of areas, such as massive arrivals of invasive species (e.g., the Pacific oyster in the Wadden Sea [5]).

Estimates of connectivity can be provided from particle transport modelling, a numerical tool in which oceanography is the indispensable input. Particle transport modelling gives substantial information on the complexity of interaction between particles and its environment. This type of model is a powerful, well-used, and adaptable method to estimate connections among marine areas, and thus, representing connectivity [6, 7]. They are desk-based and allow the simulation of a significant amount of transported particles. These tools have often been used for the transportation of small elements with little or no ability to move, which interfere with current advection. For connectivity studies, they have provided relatively accurate estimates of trajectories between locations in the oceans and the flux of individuals within trajectories. In a research context seeking for sustainability of human activities and predicting the effects of global warming, many modelling studies have examined the connectivity of marine populations, through the larval dispersal of commercial species or umbrella species (e.g., habitat formation) [6, 7]. Results from these models have contributed to understand marine connectivity and provided information for marine spatial conservation planning such as the prioritisation of zones in MPA management [8].

Across time, marine connectivity is likely to change due to the high temporal variability of the atmospheric conditions forcing surface ocean circulation [9]. The main circulation current is bound to have variation in its velocity amplitudes, geographical ranges for being wider or deeper, and directions over time, from a day-by-day basis to decades (e.g., the Gulf Stream [10]). Mesoscale circulations, such as gyres and fronts, have similar fluctuations in addition to their occurrence in the regional hydrodynamics. Circulation patterns may be accentuated and modified by climate change, to which, observations and models have shown and projected perturbations [11–13]. Temporal variability in water properties (e.g., temperature) affects ecological factors (e.g., reproductive dynamics and larval pelagic duration) and therefore, affects marine connectivity of populations, as has been shown by many biophysical modelling studies [14–16].

Knowledge of the seasonal and interannual variability of connectivity processes sheds light on the sites that have stable or intermittent connections over time and is useful for management and monitoring marine environments at middle and long term temporal scales. Connectivity estimates are presented or addressed on a temporal unit basis (e.g., comparison of connectivity between years), but the temporal variation of the connectivity over time is less represented. Only a few studies (e.g., [17–19] have conducted temporal variability approaches in the context of conservation managing plans. Estimating the variability of connectivity through time is a way to detect the sensitivity of connectivity to time, especially during strong anomalies in marine currents (i.e., years with strong events of deep-water formation, cascading, heatwaves, and storms) and during the upwelling processes. Additionally, it is also a way to identify zones where connectivity can be similar among zones or where the patterns look unique. Overall, information on the temporal variability of connectivity combined with ecological traits from selected marine species can support the policy-makers with conservation decisions. Nonetheless, it lacks of concise and adapted indicators.

In this study, we aimed to define three theoretical indicators for approaching the temporal variability of occurrence, strength, and frequency of connectivity in an area. A first calculation of the indicators was set up in semi-theoretical context using the estimates of oceanographic connectivity in the northwestern Mediterranean Sea across time obtained by numerical modelling (i.e., coupling a particle transport model with regional hydrodynamics). In addition, we investigated the impact of two relevant features impacting connectivity on the indicators estimates: the transport duration and the depth of particle sources. The proposed three indicators are thought to be ecological metrics for assessing the variability of the connectivity for zones of spatial conservation interest, and for enhancing and improving the monitoring and governance of marine resources.

## Materials and Methods

### Definition of connectivity indicators

We defined three indicators to represent the temporal variability in connectivity at a location: Occurrence, Strength, and Frequency (Table 1). Their calculations relied on the creation of links (i.e., a connection) established by at least one transported element, here a particle, between source and destination at a selected time *D* of the transport duration. Links were stored in a dataset that contained information on the Source ID, the Destination ID, the transport duration *D*, the particle release depth (binarised as near-bottom or near-surface), the particle release time, and the considered time units (month and year). We did not considered accumulation of links that were established before the time D in the calculations. Calculations of indicators did not treat ‘retention’ links differently from other links (i.e., when a source was connected to itself). Finally, for the Frequency indicator, a constant time step between releases in the particle transport was considered.

**Table 1.**
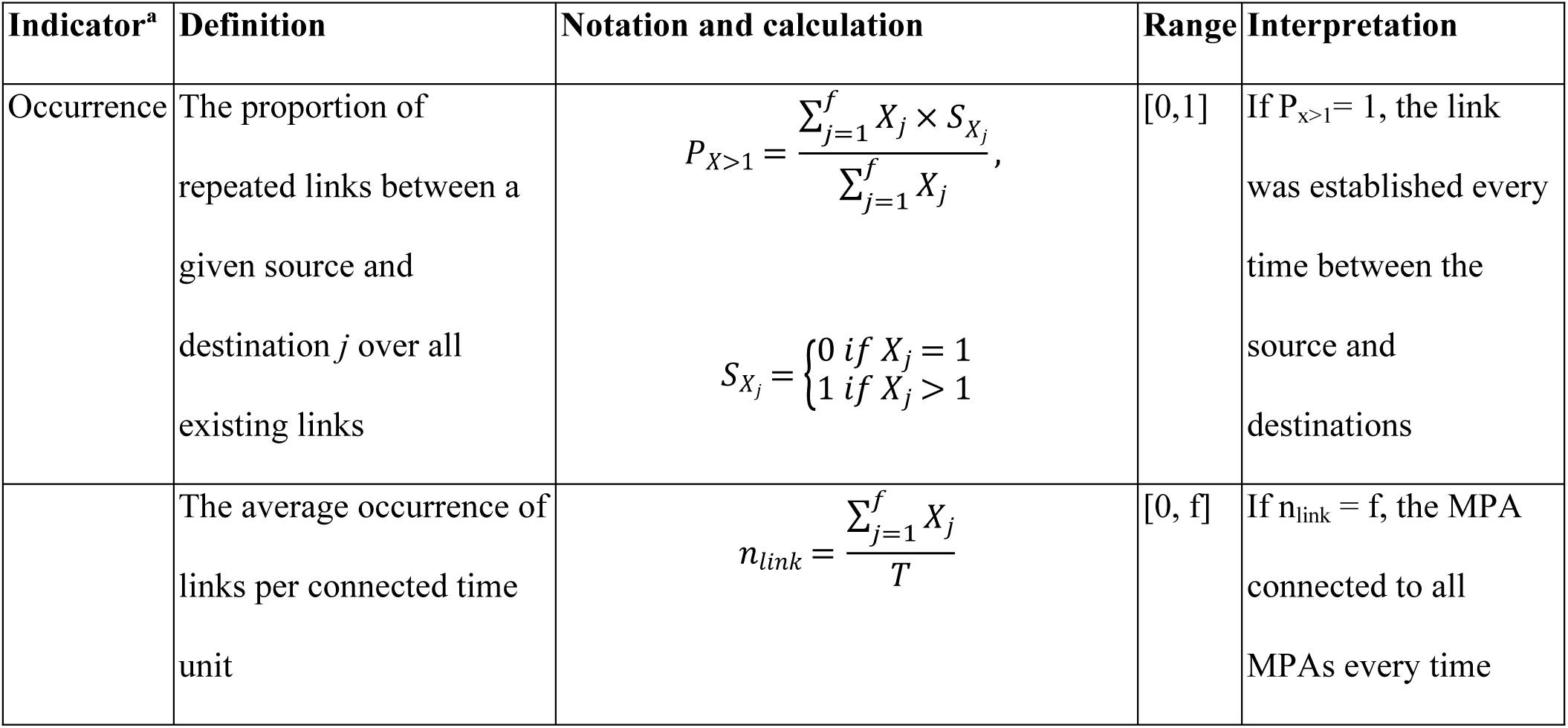
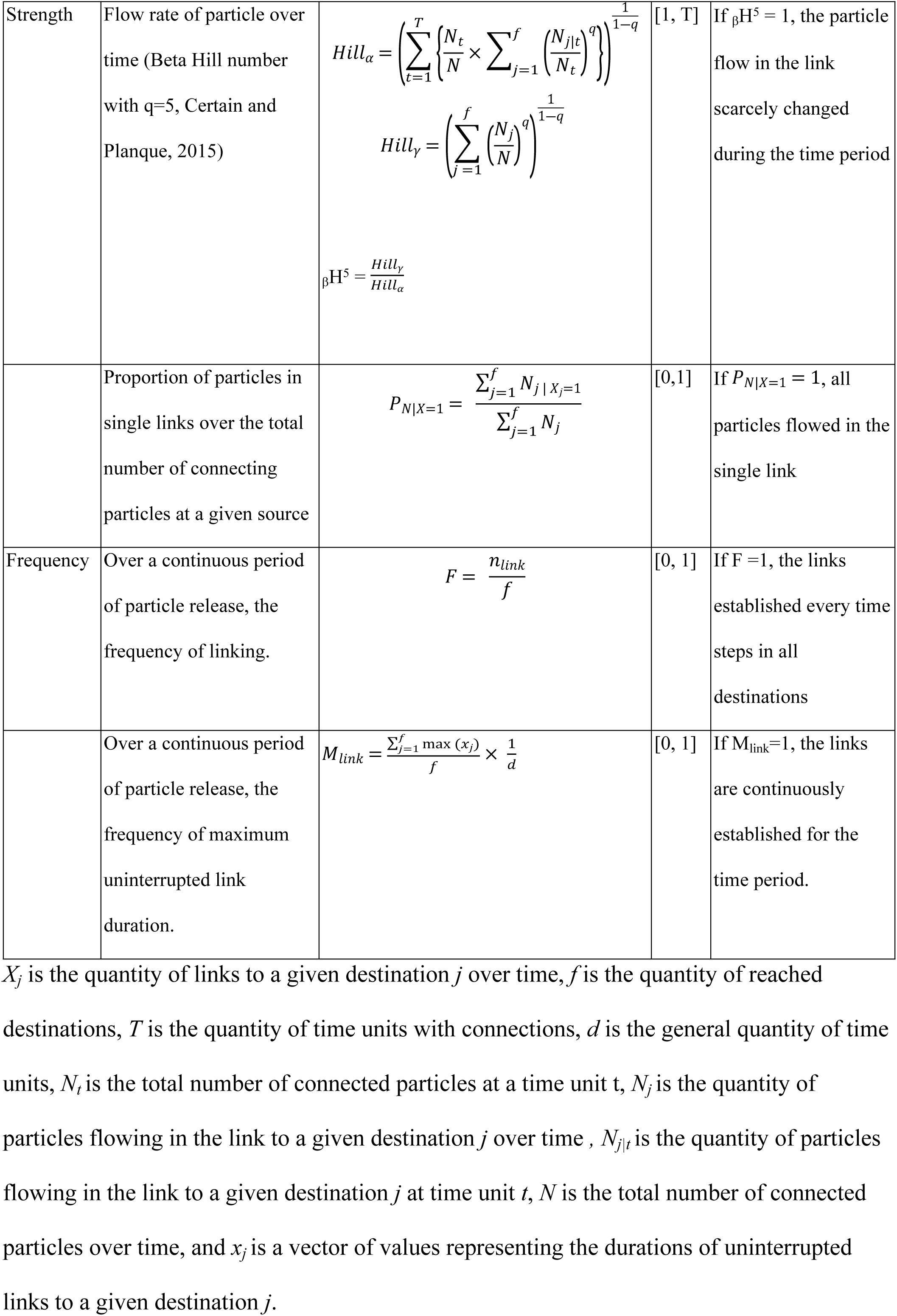
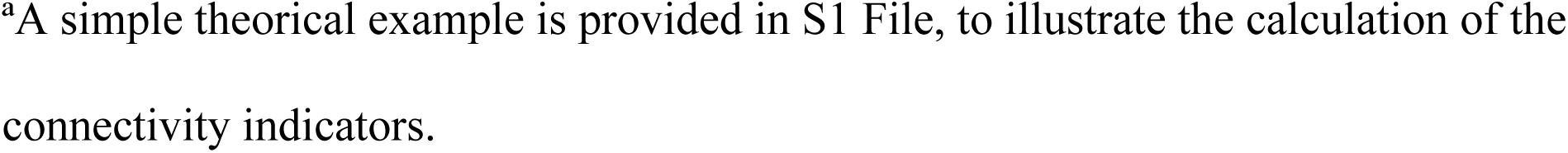
Definition of the three connectivity indicators from one source to a destination area *j*.

For the Strength connectivity indicator _β_H^5^, we adapted the application of Beta Hill number _β_H^q^, that estimates the variation of a species abundance within samples and have been found useful for marine management [20]. In our study, the particle number in one link replaced the species abundance, while the modelling year replaced the sample location. We selected the parameter *q* = 5 to emphasise the sensitivity of the Hill number to variations in links with an abundant flow.

### Biophysical modelling

#### Context of modelling

The indicators were calculated in an semi-theoretical example using the results of a Particle Transport Model (PTM) set up in the NW Mediterranean Sea (Fig 1). Such context was configured to challenge the indicators against complex ocean hydrodynamics. In the NW Mediterranean Sea, there is a well-studied southward current following the Spanish Mediterranean coastlines [21]. Mesoscale circulations introduce variability in the local circulation (e.g., gyre position, sizes, meanders along the general current), make the general circulation pattern to spatially fluctuate over time [22], and affect connectivity [14].

**Fig 1.**
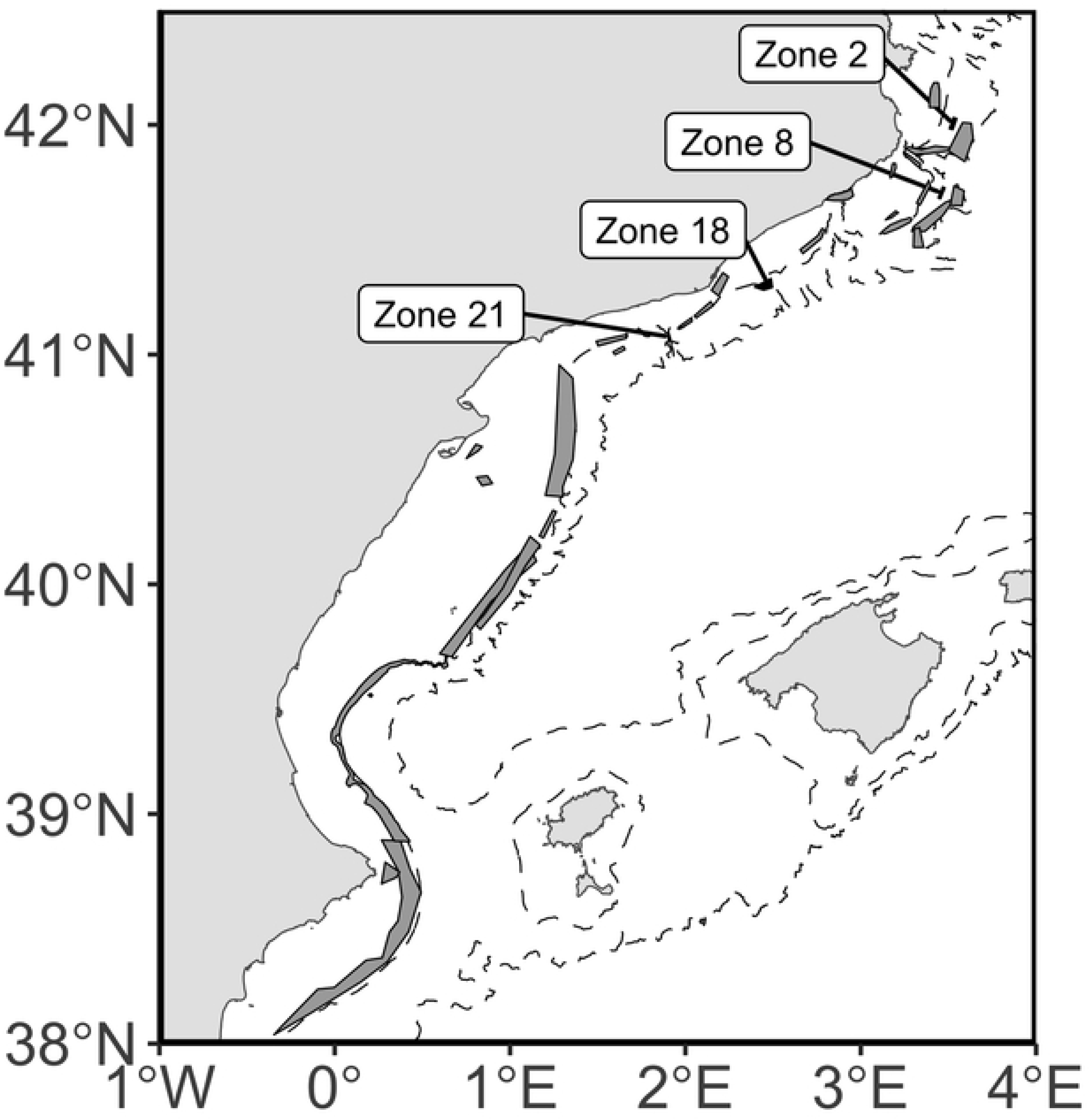
Zones of particle release and potential destinations in the NW Mediterranean Sea. The four zones with labels are the release areas of particles.

To apply the indicators that we defined (Table 1), the present case study restricted their calculations on selected areas out of 35 zones (see Fig 1) in the NW Mediterranean Sea. These 35 zones officially corresponded to marine protected areas [23, 24], but their management status was discarded from the models, and the mention of these MPAs was kept to “zones”. More particularly, four zones were selected for particle releases in the particles transport models. These four zones have similar depth distribution (300-600 m) along the continental slope and sided a well-known hydrodynamic front, having an interannual impact on larval connectivity [14].

#### Hydrodynamics

To force the advection of particles, the PTM used three-dimension velocity fields of a climatological (i.e., average circulation) and an interannual Regional Ocean Modelling Systems (ROMS [25]) run for the NW Mediterranean sea within the domain 38° N – 43.69° N, 0.65° W – 6.08° (Fig 2). The velocity fields from the hydrodynamic model are provided daily for the climatological year and between the years 2006 and 2020 on a grid with a spatial resolution around 2 km and vertically discretized over 40 sigma layers. Details on the implementation and validation of the climatological years are provided in [26, 27]. The 2006-2020 model has been forced with interannual atmospheric data provided by the Copernicus’ Climate reanalysis database (ERA-Interim from 2006 to 2016 and ERA-5 from 2016 to 2020 [28, 29]) and provided by the Cross-Calibrated Multi-Platform (version 3) from the Remote Sensing System [30]. Data were standardized to the ROMS model grid and to a time dimension of 6 hours for the air temperature, cloud cover, freshwater flux, wind components, and surface pressure and 12 hours for the shortwave and longwave radiations, and precipitation. Further details on the ROMS calibration and validation are provided in [14]

**Fig 2.**
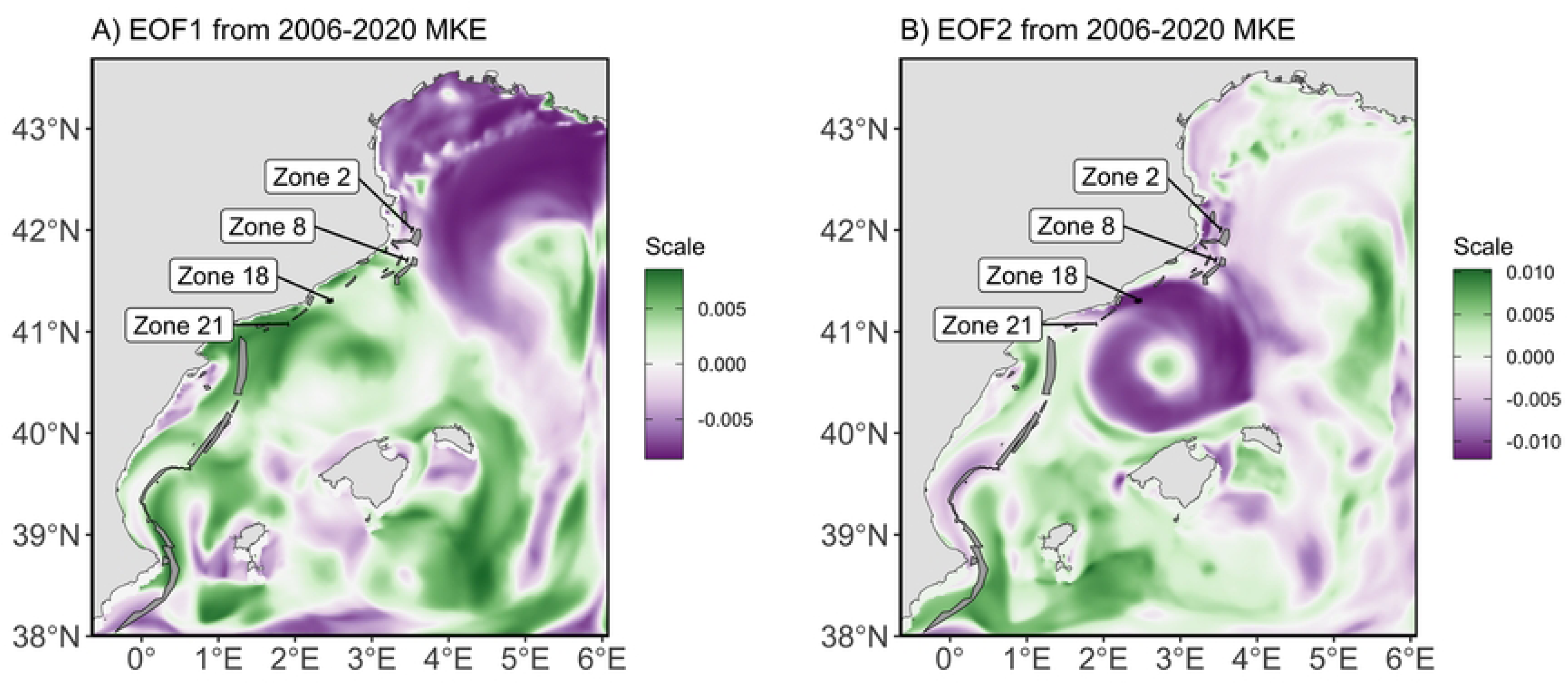
Spatial structures of the EOF1 (A) and EOF2 (B) modes of the MKE (m^2^/s^2^) during 2006-2020 in the NW Mediterranean Sea. The network of MPAs is represented by grey polygons. Labels indicate the four MPAs (2, 8, 18, and 21), used as source of particle releases.

For the implementation of the PTM with interannual variability, we used current velocity fields, whose years were selected after an Empirical Orthogonal Function (EOF) analysis on the yearly surface Mean Kinetic Energy (MKE). The analysis on the MKE revealed two main spatial structures, represented by the first two modes of the EOF analysis (explaining 38.5% of the total variance) (Fig 2). The first mode of the EOF (EOF1) represented a meander structure, which is associated with the dynamics of the Northern current, flowing NE over the Gulf of Lion. The second mode of the EOF (EOF2) illustrated an eddy (i.e., gyre) structure, which is linked to the hydrodynamics near the Balearic Sea, formed by the confluence of the Northern current over the continental shelf and a counter current formed offshore, near the Balearic Islands.

The temporal variability of the mesoscale structures allowed identifying extreme conditions in the years 2008, 2014, and 2018 among the 15 available years. The hydrodynamics of the three years were used for the runs of the particle transport model and test the performance of the indicators (Fig 3). The years 2008 and 2018 showed two local maxima in the first EOF mode, with the absolute PC values in the year 2018 being the most extreme values (0.45). The years 2008 and 2014 had respectively the lowest (-0.70) and the highest (0.43) PC values in the second EOF mode (Fig 3).

**Fig 3.**
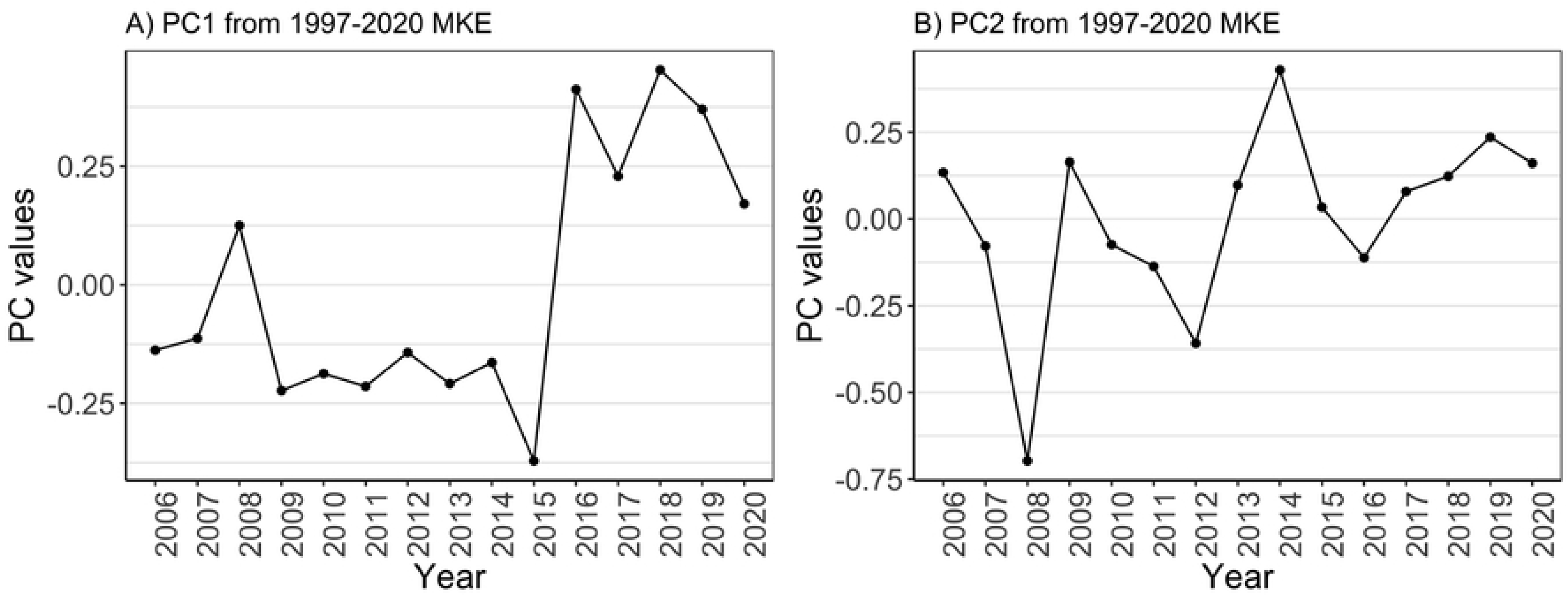
Temporal variation of MKE structures from the EOF analysis. Principal component (PC) values for EOF1 (A) and EOF2 (B) are shown for the period 2006-2020.

#### Transport modelling

Particles transport was modelled with the open source Opendrift software [31], using a Runge-Kutta 4^th^ order scheme for particle advection. Time step for trajectory calculation was set to one hour and the particle position were daily saved. If a particle hit land, the particle was deactivated and stayed at the same position until the end of the simulation.

Passive particles were randomly released within the spatial delimitation of the four selected zones 2, 8, 18, and 21 (Fig 1). The depth of the releases occurred near the surface (5 m below the surface) or near the bottom (5 m above the bottom) in a climatological year and near the surface in the selected years 2008, 2014, and 2018. The simulation of the particles’ transport started every first day of the month in the three separate years. The representativeness of the particles as early life stages of marine species has been limited when setting up the models to keep the approach as theoretical as possible. However, the transport duration lasted a maximum of 30 days, which was a value encompassing the pelagic duration of many marine species (e.g., [32]). Releases at higher frequency (e.g., every day of a month) were not implemented as we were not interested in the temporal variability of the connectivity at small time scales. A total number of 88,827 particles were released each month. This number represented more than 95% of the dispersion variability accordingly to preliminary analyses following [33] (S1 Fig.) and was distributed in the four selected zones as a pro-rata of their surface area (Zone 2 (6.4 km^2^): 13,214; Zone 8 (9.1 km^2^): 18,844; Zone 18 (23.8 km^2^): 49,110; Zone 21 (3.7 km^2^): 7,659). At the end, we ran 60 simulations and got 5,329,620 trajectories.

### Calculations of connectivity indicators

The connectivity indicators were calculated using different combinations of particle simulations to isolate effects of time variability or PTM parametrisations (Table 2). Except for the analysis of the transport duration effect on the indicators (see below), we selected the time D of 10 days for calculating the indicators. In the present study, D represented the moment a particle stopped being transported, which, in larval connectivity studies, corresponds to either the moment larvae would settle after reaching competent stages and potentially favourable habitats or the moment they would die. The settlement age is dependent to the species and to environmental contexts, hence several values of D were used in a sensitive analysis. Because our study used three discontinuous years representing extreme conditions (2008, 2014, and 2018, we used monthly outputs to compute the Frequency indicators (see Table 2).

**Table 2.** Combination of the particle transport simulations to study the time variability and model parametrisation for the indicators’ formulation.

| Perspective | Analysed effect | Indicators | Time units | Release depth | D | Total simulations |
| --- | --- | --- | --- | --- | --- | --- |
| Source | Yearly variability | Occurrence, Strength | Years: 2008, 2014, 2018 | Surface | 10 days | 36 |
|  | Monthly variability | Frequency | Months of the three years | Surface | 10 days | 12 |
|  | Transport duration | Occurrence, Strength | Months of the climatological year | Surface | 5, 10, 15, 30 days | 12 |
|  | Release depth | Occurrence, Strength | Months of the climatological year | Bottom and Surface | 10 days | 24 |
| Destination | Yearly variability | Occurrence, Strength | Years: 2008, 2014, 2018 | Surface | 10 days | 36 |
|  | Monthly<br>variability | Frequency | Months of the three<br>years | Surface | 10 days | 12 |

D: Transport duration.

In addition to focusing on the connection from a source perspective, we also calculated the indicators described in Table 1 from the reception perspective to illustrate their bidirectional use (Table 2). - The calculation remained the same, but the definitions and interpretations needed to be rewritten in terms of reception perspective at a destination zone. Connectivity indicators under the reception perspective sum up which destinations are the most and least connected to sources.

We tested the PTM parametrisation by calculating and comparing indicators for a selected transport duration D (5, 10, 15, and 30 days) and release depths (near the sea bottom and near the surface).

## Results

The connectivity between the four sources and 35 destinations are summarised in connectivity matrices (Fig 4) for the three years, an average of the three years (‘Avg’), and the climatological year (‘clim’). For D=10 days, we found that: first, for the three years, the sources did not connect with all the 35 destinations (8 zones with the IDs 26-28 and 31-35 were not connected to the sources). Second, the connectivity from the sources could be limited to a range of destinations. For example, source zones 2 and 8 were not connected to southern destinations (zones identified with an ID ≥ 17). Lastly, on a yearly basis (i.e., 2008, 2014, and 2018), the connectivity matrix also highlighted the connectivity variability per sources. For example, source zone 2 was well connected throughout the three selected years while the connectivity of source zone 21 to several zones mainly occurred in year 2014. All this diversity of connectivity temporal variability is summarised in the connectivity indicators. Based on the 3-year average, the zone 18 was connected to 22 zones, whereas this result were an outcome for the year 2014. On the contrary, results obtained from the climatological year show that there are only three connections between zone 18 and destination zones. These simple examples briefly illustrate the purpose of analysing the temporal variability of connectivity.

**Fig 4.**
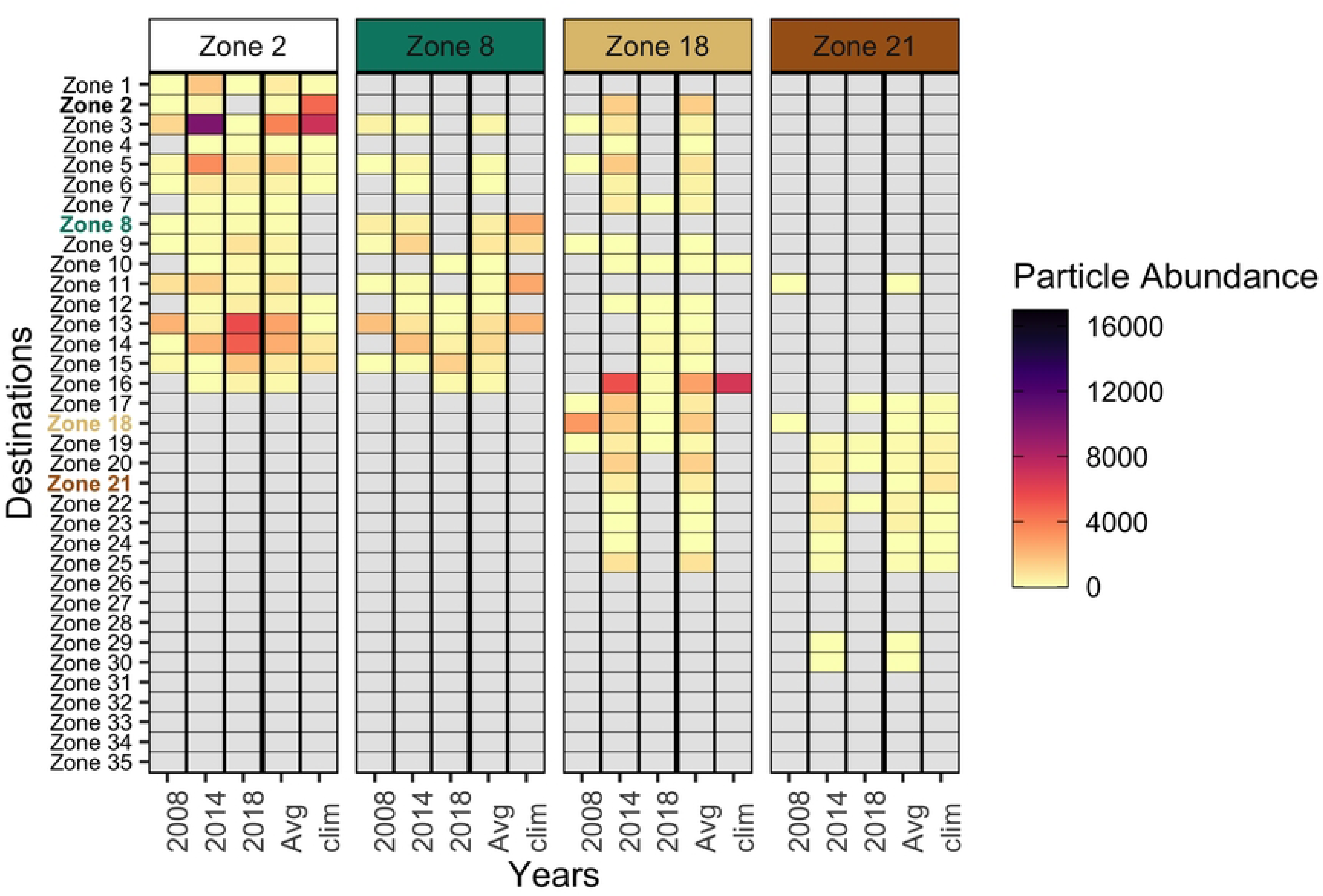
Connectivity matrix between the zone sources and destinations across three selected years, their average (‘Avg’) and the climatological year (clim’). Colour gradient represents the particle abundance flowing from a source to a destination. The grey-shaded boxes stand for zero connected particle. Zone destinations are ordered by their latitude over the NW Mediterranean Sea: Zone 1 being the northernmost and Zone 35 the southernmost.

### Indicators and time variability

#### Source perspective

The four source zones of particles presented diverse connectivity temporal variability, as shown by the scattered distribution of the source ’s indicator values (Figs 5-7).

**Fig 5.**
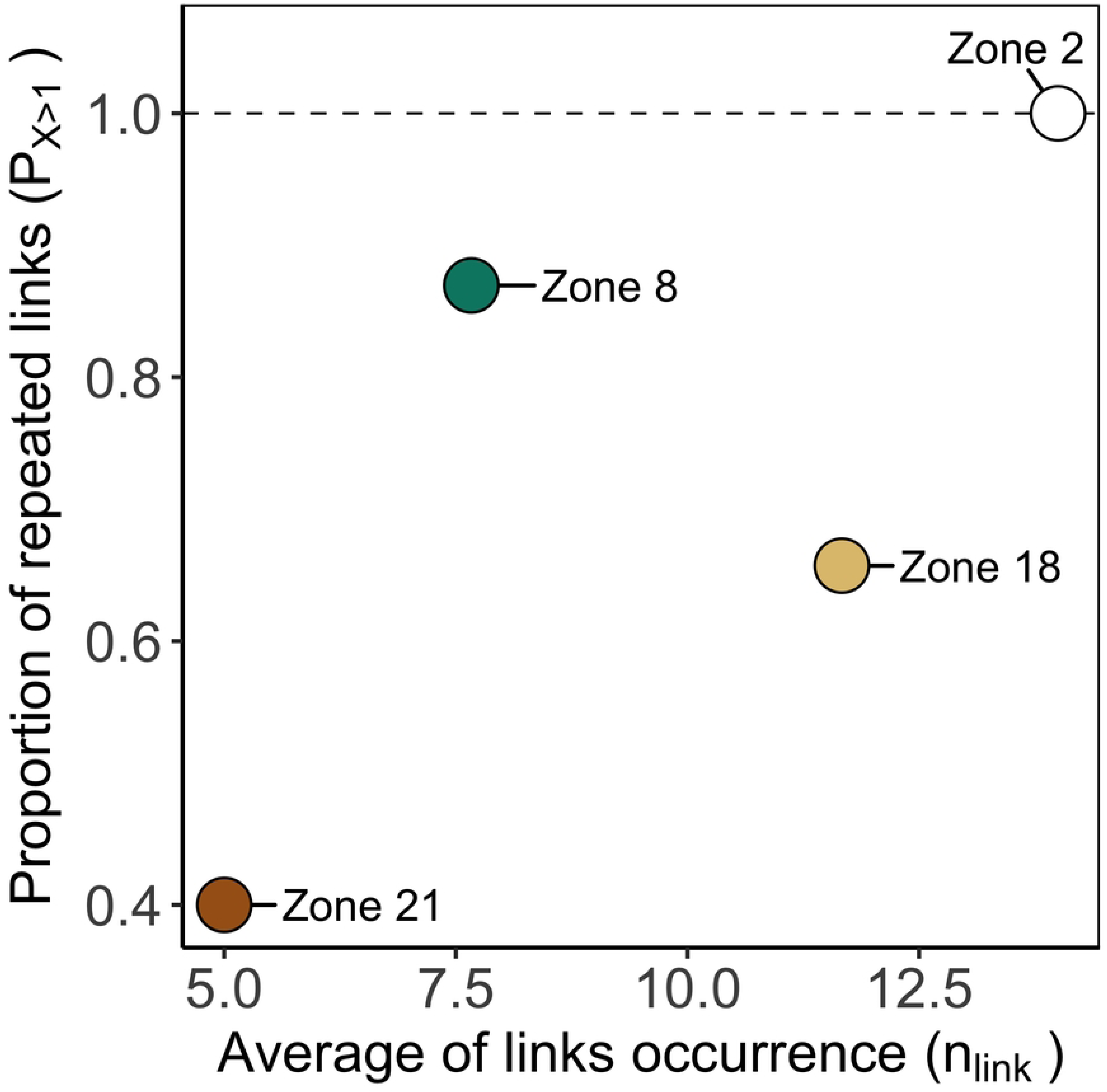
Occurrence indicators of temporal variability of connectivity according to zones as a source of particles. Source zones - Zone 2: white, Zone 8: green, Zone 18: light brown; Zone 21: brown.

**Fig 6:**
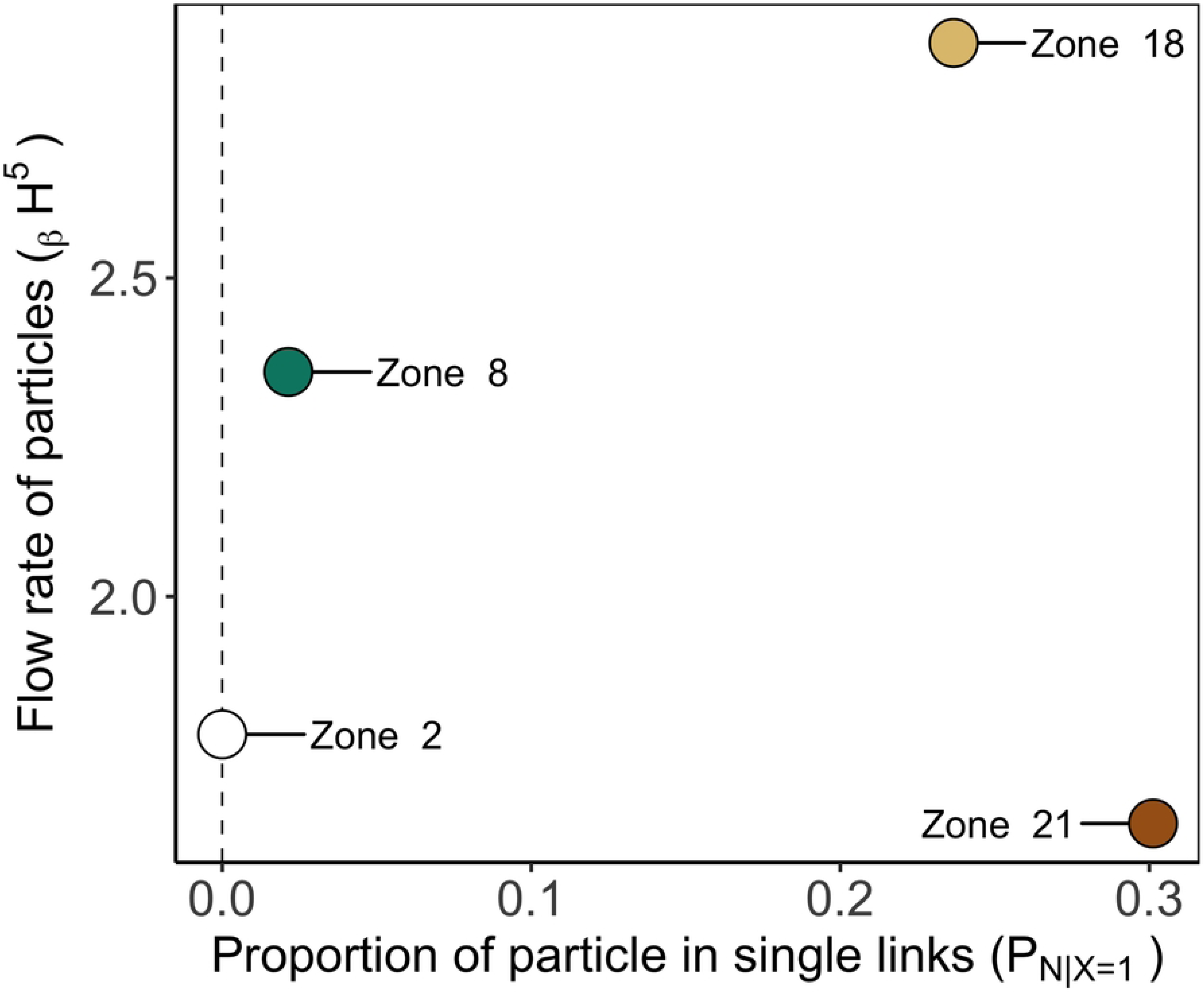
Strength indicators according to zones as a source of particles. Source zones - Zone 2: white, Zone 8: green, Zone 18: light brown; Zone 21: brown.

**Fig 7.**
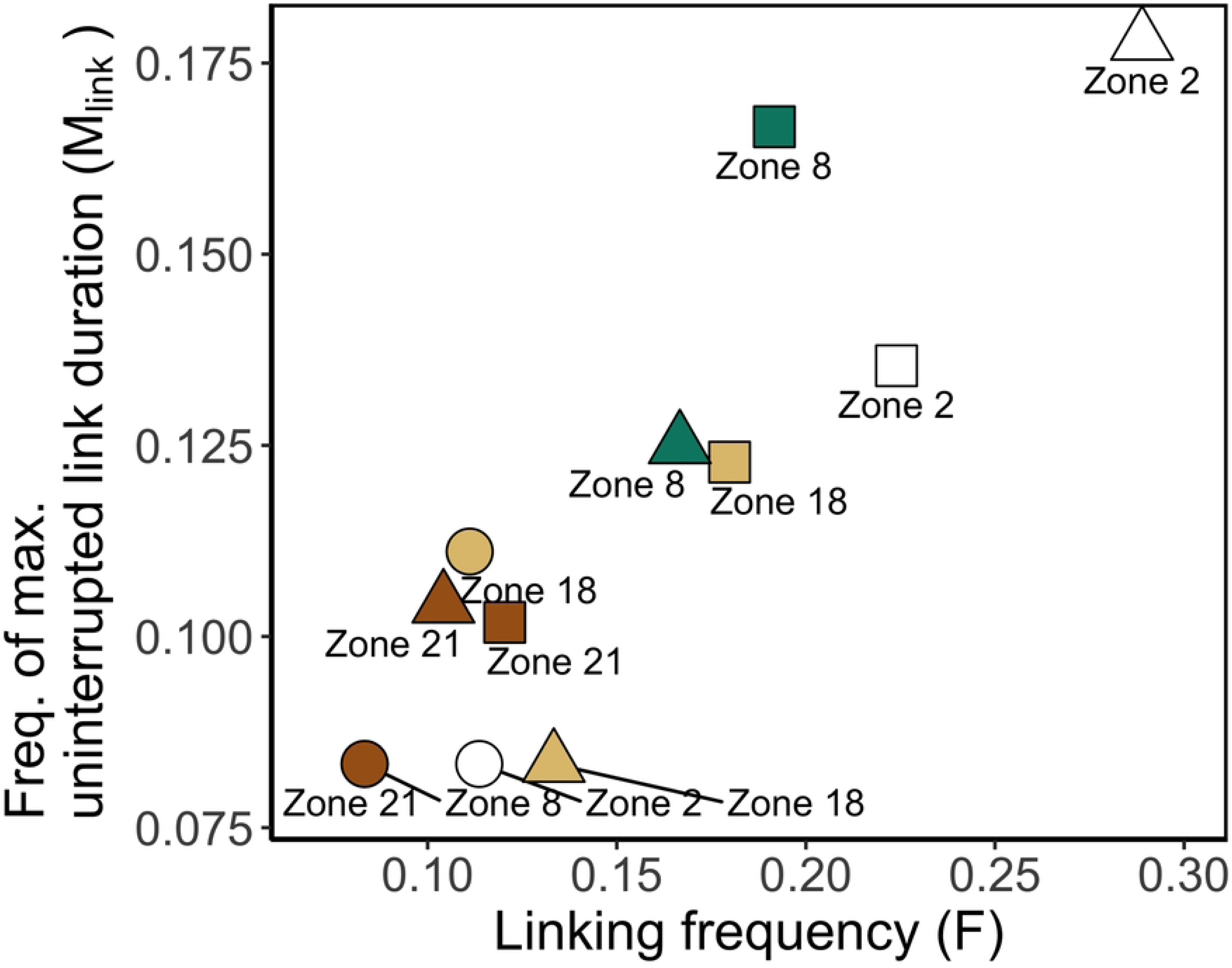
Frequency indicators according to year and zones as a source of particles. Source zones - Zone 2: white, Zone 8: green, Zone 18: light brown; Zone 21: brown. Years - 2008: round shape, 2014: square shape, 2018: triangle shape.

The Occurrence indicators (Fig 5) showed that connectivity among zones in the NW Mediterranean Sea is highly diverse. For instance, source zone 21 tended to connect a single time to a destination more often than repeatedly (*P_X>1_* = 0.4) over the three simulated years and, on average, was linked to fewer destinations (n_link_ = 5) than the other sources.

Conversely, source zone 2 was always - and at least twice - linked to the same destinations (*P_X>1_* = 1) and connected to more destinations (n_link_=14) than the other source zones. The same can be observed by analysing the connectivity matrix (see Fig 4). Source zone 21 connected with 9 destinations once, mostly in the year 2014 (6 single links) and source zone 2 connected at least twice with 16 destinations (thrice with 10 destinations). These Occurrence indicators alone already highlight the relevance of zone 2 position for likely creating persistent links with others. On the spatial dimension, zone 2 is one of the northernmost sites of the studied area and is geographically close to 11 zones in a radius of 50 km (see Fig 1). Its proximity to several zones can help to have a high *n_link_*. However, the proximity of other zones does not necessarily imply *n_link_*would always be high. This statement is illustrated by the less connected source zone 8 (n_link_ = 7.67), which was geographically close to 14 zones within a radius of 50 km, while it was located 30 km south of source zone 2.

The Strength indicators (Fig 6) revealed the subtleties across time occurring in the established links, and this statement also applies for the links occurring just once. Source zone 21 can be considered a source with limited flow of particles among the source zones because connections were rare and lowly impactful over the selected years. Indeed, the flow of particles through the single links P_N | X= 1_ from source zone 21 represented 30% of the particle flow. Besides, its flow of particles towards the other destinations was relatively constant (_β_H^5^ close to 1). These statements were also visible within the connectivity matrix (Fig 4). Less than 200 particles were transported through the links from source zone 21. To a lesser extent, source zone 18 also had low-impactful single links but had +more variations of particle flows through the repeated links (_β_H^5^ close to 3). For source zone 2, Beta Hill number highlighted that the flow of particle relatively and mildly fluctuates between the three selected years. This would mean that the position of zone 2 is not only strategic to connect with several destinations several times (i.e., high n_link_), but also to sustain a consistent connectivity flow across time, making that zone a potential super spreader site.

The Frequency indicators could discern the monthly variability of the connectivity in the years 2008, 2014, and 2018 and among the four sources (Fig 7). On the first hand, the monthly connectivity in the years 2008 and 2014 had opposite patterns. The low values of the indicators in the year 2008 indicated a less frequent establishment of links along the 12 month period (F ∼ 0.1) and a lower probability of consecutive connections (M_Link_ = 0.087) than for the other years. It was the opposite for the year 2014: links lasted a maximum of 1.5 months (M_Link_ = 0.13) and occurred 2.1 times in 12 months (F ∼ 0.18). Taken separately (S2 Fig.), the matrices of connectivity also represent these distinct frequency features of connectivity. On the other hand, the Frequency indicators revealed that source zones have relatively comparable variation of connectivity through 12 months. Indeed, in the three studied years, the Frequency indicators of source zones 18 and 21 were generally lower (F < and M_Link_ < ) in comparison to the source zones 2 and 8 (F > and M_Link_ >).

#### Reception perspective

An alternative perspective was to consider the reception of particles at the zones of interest (i.e., 2, 8, 18, and 21), in other words, as destination zones (Fig 8). We found that all the zones, except zone 21, are connected at least once per year (n_link_ ≥ 1) and are bound to receive a similar flow of particles every time they are connected (_β_H^5^ ∈ [1 - 1.3]). Particular cases of connectivity variability were also highlighted like the high proportion of particles channelled towards zones 2 and 21 in single links (P_N | X= 1_ > 0.8) or like the low Frequency indicators for zone 21 (F ∼ 0.09 and M_Link_ ∼ 0.08). These Frequency indicators and the information from a source perspective confirmed that the connectivity impact of zone 21 among the four zones was limited as a source and a destination.

**Fig 8.**
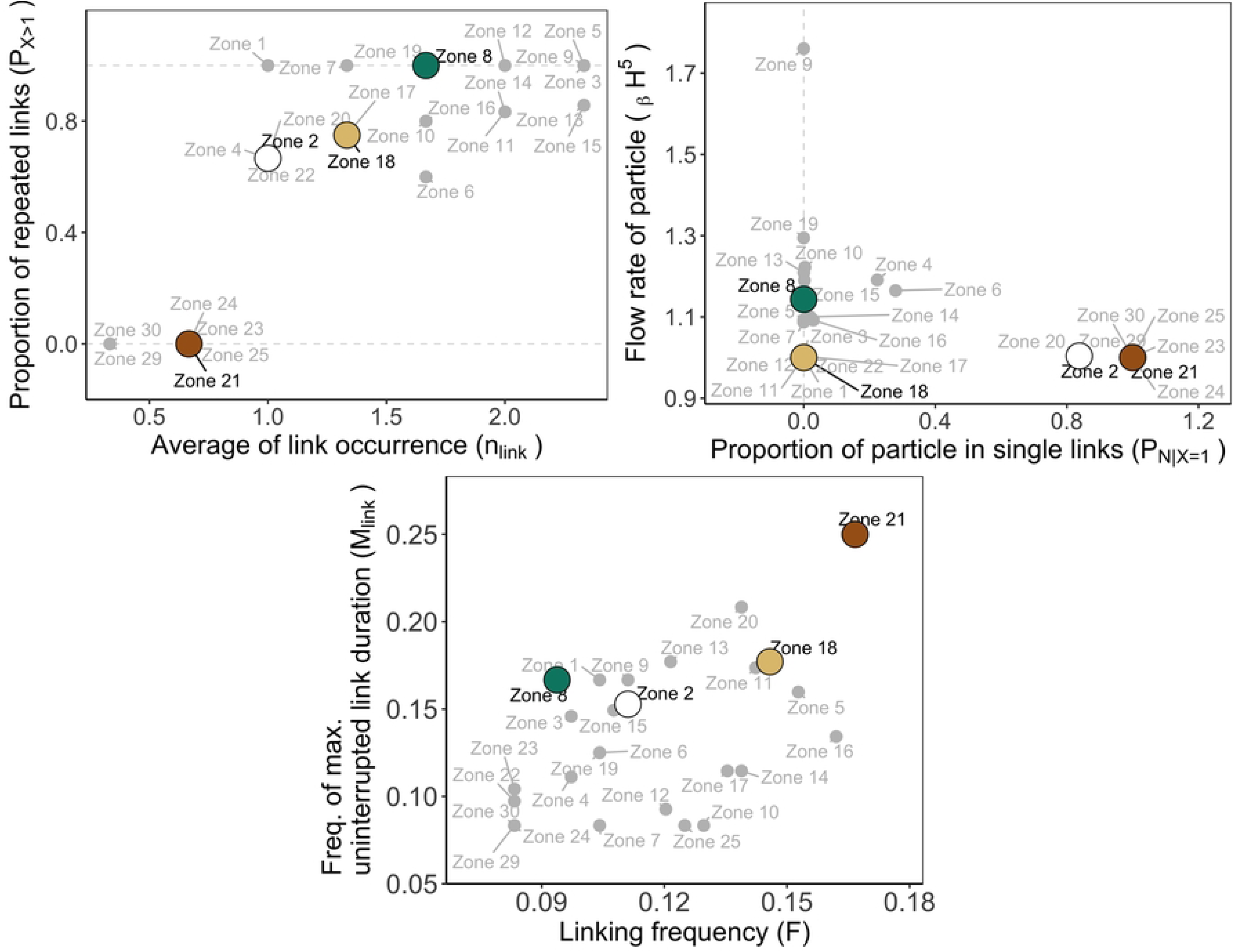
Occurrence (A), Strength (B), and Frequency (C) indicators according to zones as a destination of particles. Source zones - Zone 2 white, Zone 8: green, Zone 18: light brown; Zone 21: brown. Other zones: grey.

Recontextualising the four studied zones within all existing zones (grey labels and points in Fig 8) allowed us to estimate clusters of zones with similar connectivity indicators. This showed that indicators reached much higher values when considering the 35 zones than for the four studied sources: nine destinations had a *n_link_* ≥ 2 and one destination (i.e., zone 9) had _β_H^5^ > 1.7. As for clustering zones together based on these indicadors, we could associate the destinations 2 and 21 to six other zones (20, 23-25, and 29-30 in Fig 8.B) for their similar Strength indicator (P_N | X= 1_ > 0.8 and _β_H^5^ ∼1.1) and group the destination zone 21 with five other zones (23-25, 29-30 in Fig 8.A) based on their Occurrence indicator (n_link_ < 1 and P_X>1_ = 0).

### Sensitivity of the indicators to model parametrisation

#### Release depths

The depth of releases impacted the indicators in three different ways (Fig 9). First, null or minor changes in the connectivity indicators across time and sources (e.g., n_link_) were detected in the indicators. The Occurrence indicator n_link_ shows that the number of links is likely to stay the same if the dispersal occurred near bottom or near surface (e.g., for source zone 18, n_link_ = 1.29 with surface simulations and n_link_ = 1.75 with bottom simulations). It is worth to highlight that although n_link_ changes were small with the depth of the simulation, the destinations of particles could be modified. For example, source zone 18 connected to zones 21 and 22 with bottom simulations and to zones 16 and 18 with surface simulation (S3 Fig.).

**Fig 9.**
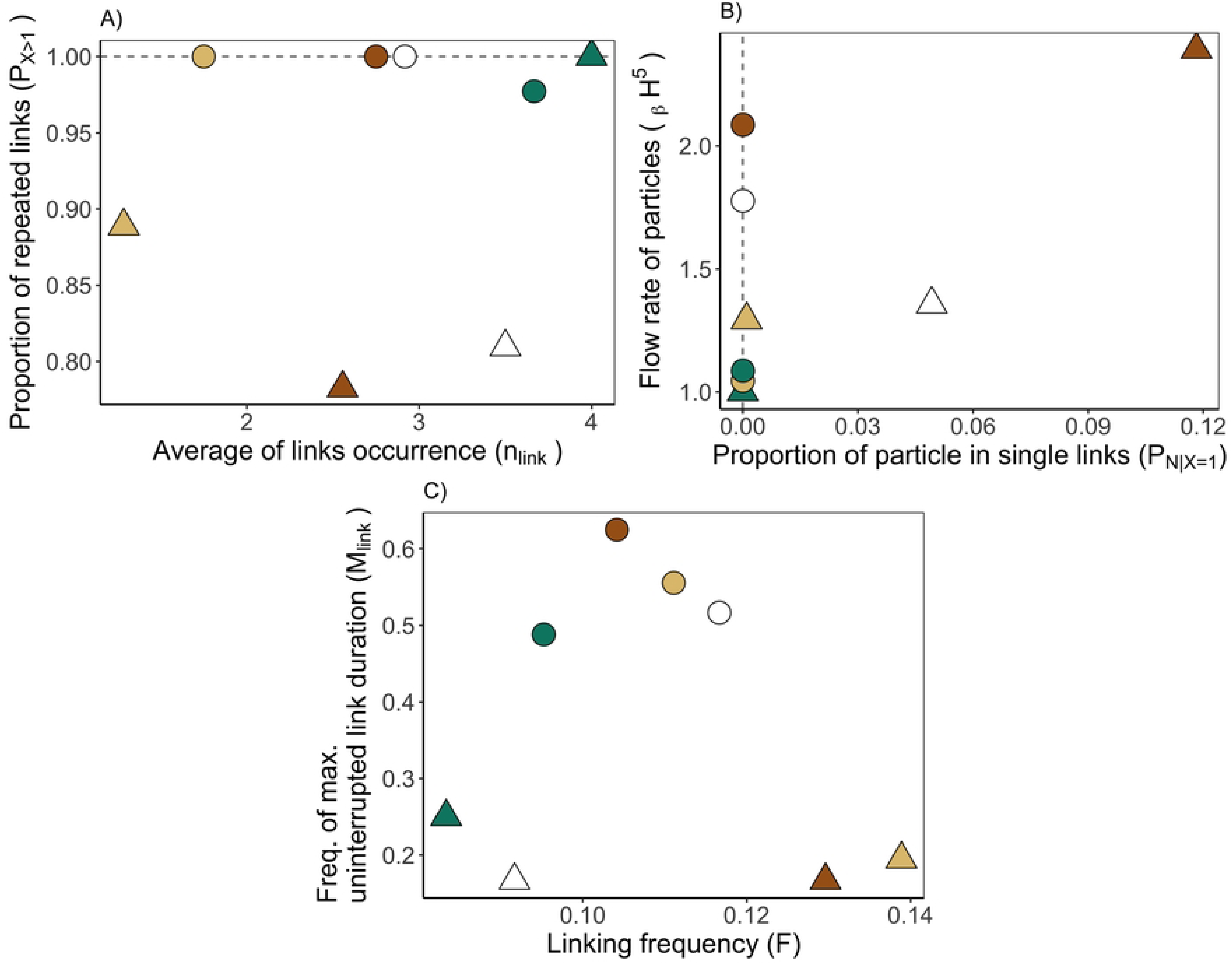
Occurrence (A), Strength (B), and Frequency (C) indicators across 12 months of a climatological year according to source zones and release depths. Source zones - Zone 2: white, Zone 8: green, Zone 18: light brown; Zone 21: brown. Release depth - near the bottom (round shape) and near the surface (triangle shape).

Second, the indicators for a few source zones was modified (e.g., _β_H^5^ and *P_N|X=1_* for zones 2 and 21 in Fig 9.B). For example, the releases made near-bottom decreased and increased _β_H^5^ in source zones 2 and 21, respectively, in comparison to releases made near-surface. Finally, the depth of release marked a clear dichotomy in indicators for most (if not all) zones (e.g., P_X > 1_ and M_link_ in Figs 9A and 9C). The probability that the source had a repeated link to a destination *P_X>1_* decreased by 0.17 for releases near the bottom (*P_X>1_* = 1.00) in comparison to releases near the surface (*P_X>1_* = 0.83 on average) for all sources except source zone 21. The dichotomy was also well seen in the Frequency indicator M_Link_, with longer continuous connections for particle released near the bottom (M_Link_= 0.55 on average) than near the surface (M_Link_= 0.19 on average).

#### Transport duration

Analysing the sensitivity of the connectivity indicators to the transport duration through the selection of times D revealed that there are multiple possibilities of connectivity variations depending on the source (Fig 10). The indicator value increased/decreased with increasing D from 5 to 30 days. For example, P_X>1_ from 0.95 to 0.78 in source zone 2 and _β_H^5^ from 1.001 to 1.052 in source zone18. Some indicators had small or big variations in their values without linear relationship to the transport duration, e.g., source zone 2 with M_link_ ∈ [0.12, 0.24] and source zone 21 with P_X>1_ ∈ [0.85, 1.0]. The zones could also be grouped according to relatively low and high amplitudes of indicators variations, suggesting that there was a different impact of the transport duration based on the source zone, which could be additionally associated with their geographical position in the NW Mediterranean Sea. For example, the indicator variations were null for the P_X>1_ and P_N|X=1_ of source zone 8 (standard deviation = 0) while standard deviation of P_X>1_ over all transport durations was between 0.05 and 0.07 for the other sources. The M_link_ was lower (i.e., 0.13 on average) for southern source zones 18 and 21 than for northern source zones 2 and 8 (i.e., 0.16 on average).

**Fig 10.**
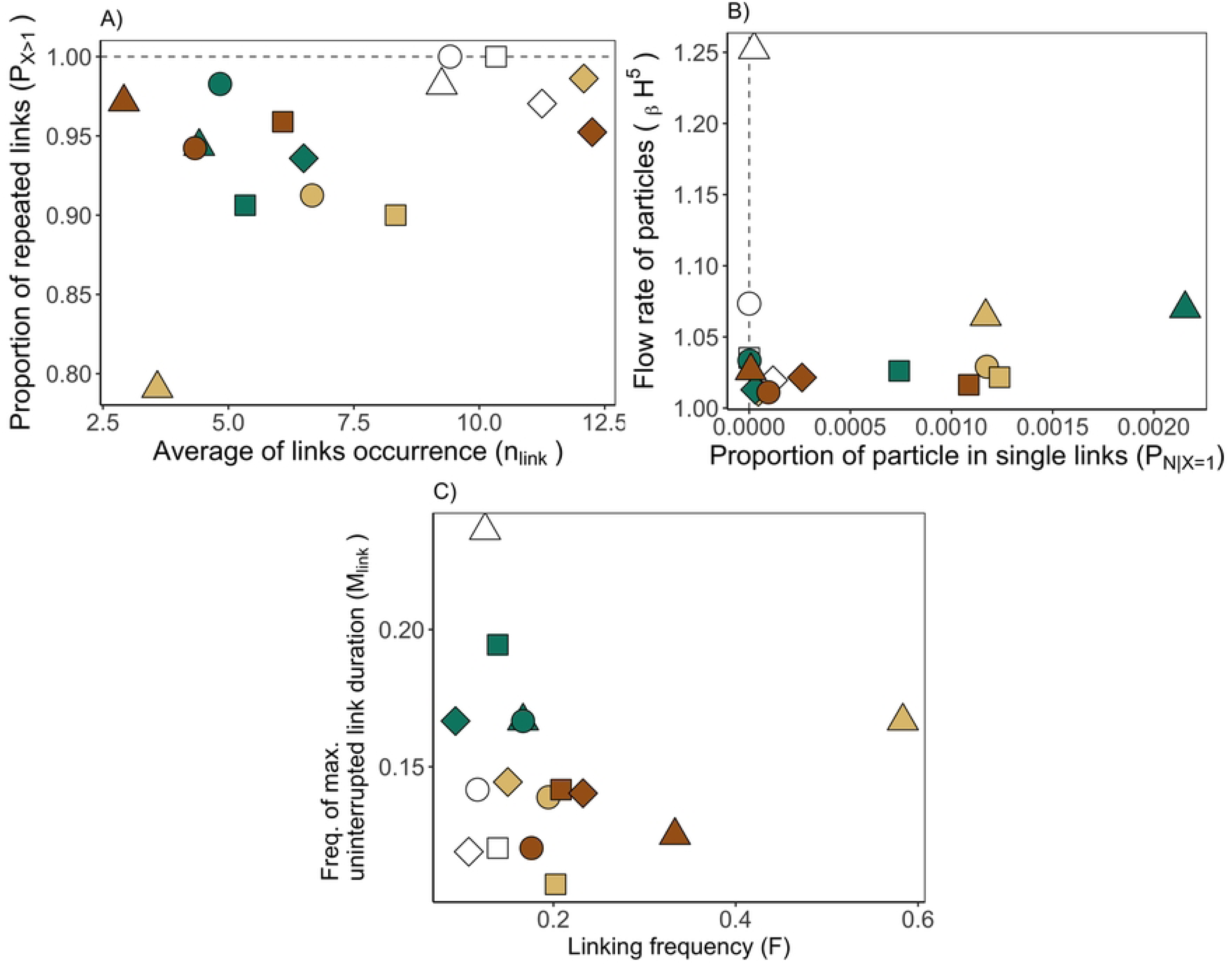
Occurrence (A), Strength (B), and Frequency (C) indicators according to transport durations and zones as source of particles. Transport durations - 5 days: triangle; 10 days, round; 15 days: square, 30 days: diamond. Source zones - Zone 2: white, Zone 8: green, Zone 18: light brown; Zone 21: brown).

## Discussion

The proposed indicators allowed us to characterise the variability of connectivity from sources and destinations and contributed to identifying zones for spatial conservation management that have potentially high oceanographic connectivity through time.

### Temporal variability of the connectivity

The temporal variability of the connectivity was influenced by the spatial distribution of zones along the NW Mediterranean margin. On the one hand, their geographical distribution in the area played a role in the variability of connectivity due to their exposure to the dominant water circulation, as often observed in other regions [34, 35]. The formation of multiple connections over time, defined by the Occurrence indicator *P_X>1_* and its inverse (i.e., 1-*P_X>1_*, when a source zone tends to connect a single time to another zone) highlighted how some zones were more exposed to the temporal variability of currents than other zones. For example, single connections may occur when particles are advected in velocity fields (by their directions or amplitudes) established under abnormal circumstances (e.g., under the influence of a cyclone [36]). A low *P_X>1_* implied that the water circulation would normally isolate one zone from the rest and that, for some years, anomalies in the circulation pattern would suspend this situation and allow connectivity (e.g., results on zone 21). In our case, the anomaly was due to a strong southward current in 2014 contrary to the circular current observed in the years 2008 and 2018. Such results recall the importance of having hydrodynamic models that well represent the mesoscale structures of the circulation. A fine resolution of the model is a prerequisite for a good representation of connectivity [37], especially when larval dispersal occurred in coastal waters. Our study was less sensitive to the hydrodynamic model resolution of 2 km because it was conducted in open waters and for zones above the continental slope. However, if a finer model were available, the differences in connectivity indicators between zones would probably be more pronounced. On the other hand, the distance between the zones (i.e., the spacing between sources and destinations) as a mild impact on the connectivity temporal variability: a network of close zones did not necessarily imply a higher occurrence of connections (e.g., low *n_link_* for source zone 8) in our theoretical approach. Spacing is a relevant criterion for defining zones as MPAs, following proposed recommendations [38, 39]. Yet, if the geographical distribution of spatially close zones always overlaps with interannually variable mesoscale structures, as in zone 8, connectivity and its variability may be restricted and less relevant for MPA performance. From this theoretical finding, we suggest that the persistence of local hydrodynamics should also be investigated when designing spatially-close MPAs.

The present indicators can be used to characterise zones on the basis of their success in establishing connections over the temporal variability of external conditions. For instance, a source zone with numerous connections (high *n_link_*) that are repeated over a unit of time (high *F*) can be defined as a ‘persistent’ super spreader, using the terminology of graph theory networks. If it has only one repeated connection (low *n_link_*) to a destination, it can be defined as an ‘persistent’ restricted spreader. In terms of the reception perspective, a destination zone that maintained several connections (high *n_link_*) frome time to time (low *F*) can be called an ‘occasional’ super-receiver. This classification of zones based on characteristics of temporal connectivity variability has a practical outcome for stakeholders’ understandings, especially when designing a network of MPAs whose implementation is long term. These indicators enhanced the need for further analysis of certain zones to understand whether they are relevant and essential for regional connectivity. In fact, the indicators could reveal unusual connectivity for certain zones (e.g., zone 18) where it might be strategic or not to make conservation decisions. For instance, would a protected area that tends to be connected only once and by a few particles to a non-isolated MPA destination (i.e., one that is already well connected to other MPA sources) be a priority for implementation or modification? The answer may be complex, as the impact of the presence of an MPA on connectivity in an MPA network may be quite significant [40]. Calculating the temporal variability of connectivity can provide another layer of criteria to inform management decisions when previous assessments are inconclusive.

### Connectivity indicators with ecological implications

The calculations of the indicators were based on a semi-theoretical approach and have yet to be applied to larval transport studies with specific species characterisations. The PTM parametrisations approximated key ecological traits of early-life stages of marine species, affecting connectivity: 1) depth of particle release as the spawning depth of demersal and pelagic species and also surfacing vertical larval behaviour, and 2) transport duration as the pelagic life duration (PLD) of early life stages (including the different forms of the pelagic stages, such as eggs or larvae). The sensitivity of the indicators to the parameters showed interesting results for using indicators to compare the connectivity of marine species and to understand the impact of global warming on connectivity. Firstly, the indicators are likely to be helpful in showing when early life stages with short or long PLD, or with a benthic or epipelagic habitat would be affected in an environment with temporal variability. The expected results would be in line with the many existing studies addressing the issue of ecology in connectivity [32, 41,42]. In addition, these indicators shall assess whether a zone allows connectivity for multiple species or for a single species once ecological traits are included in the transport models. It implies divergent or convergent connectivity patterns as function of the species life habits [42, 43]. This would permit to understand if the MPA network is ecologically coherent [44], meaning that it is adequately designed for various groups of species. Secondly, by measuring and comparing connectivity indicators over two periods separated by a known regime shift, the temporal variability of the connectivity due to climate change can be estimated. In the NW Mediterranean Sea, an increase in the water temperature has been observed over the last two decades [45] and correlated with various changes in marine community in the Mediterranean region [46–48]. In the transport of early life stages, the time spent in the pelagic waters could shorten or lengthen according to species-dependent metabolic and behavioural traits associated with water temperature [49]. Consequently, connectivity is likely to change over time.

Further work on the calculation of indicators should help to improve the measurement of the temporal connectivity variability when including the ecological dimension. The context of the study did not allow further investigation of more connectivity indicators but their development and calculation would be welcome. Firstly, we lacked information on the temporal quality of connectivity, which can nevertheless be approached within the calculation of the Strength indicator. The quality of the connectivity should take into account the loss of particles before, during and after the transport, and is strongly, if not exclusively, related to the intrinsic ecology of the species (e.g., spawning success, mortality, growth, or reproduction rates). These ecological traits are essential information for determining with greater precision the temporal connectivity variability of a given set of populations in a geographic network of MPAs [50]. Second, a useful Occurrence indicator would be to indicate whether an MPA is receiving and sourcing dispersing particles. This metric would make it possible to understand whether an MPA is receiving more/equal/less larvae than it is delivering to other MPAs or whether it is acting as a recurring stepping stone unit over the course of several years [51]. In fact, the methodology of the graph-theoretic approach [52], already used to design and evalute MPA networks [40, 51], should help to develop indicator calculations that focus on the temporal variability of connectivity. Third, the calculation of another Frequency indicator should help to determine the patterns of seasonality in the course of several years and provide meaningful information for the management of temporary MPA closures. At the species level, connectivity can be episodic and seasonal, since marine species have different spawning strategies and life cycles, resulting in peaks of connectivity at certain times of the year. In particular, many species release individuals (i.e., eggs, larvae, spores, propagules) either at a specific time of the year (e.g., winter in the case of *Nephrops norvegicus* in the NW Mediterranean Sea [53]), relatively frequently throughout the year [54] like the European hake *Merluccius merluccius* in the NW Mediterranean Sea [55].

### Indicators for conservation planning and assessment

The purpose of defining indicators of temporal connectivity variability is, in part, to support spatial conservation planning. Studies have shown that an optimal MPA network can be estimated from an analysis over a relatively short period of time (i.e., 10 years [56, 57]) and that MPA effects are expected to be seen after a few years [58]. Such findings suggest that: *i)* interannual monitoring and assessment of MPAs should be considered, *ii*) this optimal period of time should be searched for the MPA network in the NW Mediterranean Sea, and *iii*) the usefulness of the presented connectivity indicators should be questioned by calculating them over the selected optimal time period.

In our study, the zones were 35 deep-sea MPAs in the NW Mediterranean Sea, established for the recovery and conservation of target species of commercial fisheries (i.e., the European hake *Merluccius merluccius*, the Norway lobster *Nephrops norvegicus*, the blue and red shrimp *Aristeus antennatus*) and deep-sea bioengineers (e.g., soft-bodied cold-water corals and gorgonians). Connectivity has not yet been assessed in these MPAs, but monitoring data have begun to be collected and processed [59–60]. Calculation of the indicators of temporal variability in connectivity in this network, using a setup of ecological traits in the PTM, would provide benchmarks for MPA assessment and understandings of long-term connectivity in deep-water continental margins. These indicators appear to be a good and robust way to compare the connectivity variability of MPAs, although they do not pretend to replace other existing methods of connectivity analysis, e.g., graph theoretical indicator, connectivity matrices, connectivity portfolio effect [61]. In fact, we still need matrices to understand and quantify what is observed with the indicators. Far beyond their use for comparison, indicators can be helpful to redesign MPA networks, adapting their structure to locations with interesting long-term connectivity patterns, and thus, improving the conservation of marine ecosystems. Redesigning MPAs with PTM requires a computational effort, but it helps to optimise the connectivity in a geographic network [62]. As long as spatial positions are stored over time, connectivity indicators can be calculated on datasets obtained from other methods involved in the movement tracking, such as Lagrangian drifters (e.g. Argos buoys), which would enhance the aspect of oceanographic connectivity; as well as mark-recaptured tagging of animals, which could help understand the dispersal of the species to different zones for their ecological needs (e.g., feeding, foraging) over years. The application of the indicators is also open to different types of marine spatial management planning, such as the spread of marine invasive species and recruitment to fishing grounds.

## Conclusions

Three indicators characterising connectivity have been defined and applied in 35 zones distributed in the NW Mediterranean Sea. These indicators are based on the temporal variability in connectivity established among zones of interest (i.e., MPAs) through particle tracking modelling. They encompass the notion of connectivity Occurrence, Strength and Frequency among zones. These indicators would be relevant for decision-makers during the design process of MPAs or when reporting on their efficiency. Applying these indicators should provide and underline the similarities and differences in terms of connectivity among zones, hence refining a location appropriateness among other candidate zones to management planning. Additionally, a likely use of the indicator comparison is to estimate the capacity of the zone to allow connectivity for species with different ecological traits and the connectivity variation under climate change scenarios (e.g., increasing water temperature). Overall, the indicators are a summary of connectivity variability when multiple time steps are considered in a study. We consider that these indicators are applicable irrespectively to the studied regions (here the NW Mediterranean Sea) and to the connected zones (here MPAs), and more importantly, relevant for assessing the MPA network coherency and prioritising management efforts on some MPAs of the network.

## Acknowledgements

We thank Dr J. Solé for providing hydrodynamic datasets, Dr J. Yearsley for the help on the elaboration of a connectivity indicator, and Dr C. Artana for providing helpful papers for the discussion on hydrodynamic changes under the global warming.

## Supporting Information

**S1 File. Theoretical example for calculating Connectivity indicators**

**S1 Fig. Fraction of unexplained variance (FUV) in the particle dispersal according to the number of released particles based on Simons et al. (2013) methodology.** The dashed grey lines indicate the threshold of 0.05, below which, the number of particles represents 95% of the particle dispersal variability. The threshold is reached for 85,000 particles. Red error bars show the standard deviation of FUV around the mean.

**S2 Fig. Monthly connectivity occurrence (X-axis) from a source zone to a destination zone for the years A) 2008, B) 2014, and C) 2018.**

**S3 Fig. Occurrence and flow of connectivity over months from a source zone (facets) to a destination zone for the particle release A) near bottom and B) near surface.**

